# Kynurenine pathway in post-mortem prefrontal cortex and cerebellum in schizophrenia: relationship with monoamines and symptomatology

**DOI:** 10.1101/2021.02.22.432214

**Authors:** Amira Ben Afia, Èlia Vila, Karina S. MacDowell, Aida Ormazabal, Juan Carlos Leza, Josep Maria Haro, Rafael Artuch, Belén Ramos, Borja Garcia-Bueno

## Abstract

**Background:** the cortico-cerebellar-thalamic-cortical circuit has been implicated in the emergence of psychotic symptoms in schizophrenia (SZ). The kynurenine pathway (KP) has been linked to alterations in glutamatergic and monoaminergic neurotransmission and to SZ symptomatology through the production of the metabolites quinolinic acid (QA) and kynurenic acid (KYNA).

**Methods:** this work describes alterations in KP in the post-mortem prefrontal cortex (PFC) and cerebellum (CB) of 15 chronic SZ patients and 14 control subjects in PFC and 13 control subjects in CB using immunoblot for protein levels and ELISA for interleukins and QA and KYNA determinations. Monoamine metabolites were analysed by High Performance Liquid Chromatography and SZ symptomatology was assessed by Positive and Negative Syndrome Scale (PANSS). The association of KP with inflammatory mediators, monoamine metabolism and SZ symptomatology was explored.

**Results:** in the PFC, the presence of the anti-inflammatory cytokine IL-10 together with IDO2 and KATII enzymes decreased in SZ, while TDO and KMO enzymes expression increased. A network interaction analysis showed that in the PFC IL-10 was coupled to the QA branch of the kynurenine pathway (TDO-KMO-QA), whereas IL-10 associated with KMO in CB. KYNA in the CB inversely correlated with negative and general PANSS psychopathology. Although there were no changes in monoamine metabolites content in the PFC in SZ, a network interaction analysis showed associations between dopamine and methoxyhydroxyphenylglycol degradation metabolite. Direct correlations were found between general PANSS psychopathology and the serotonin degradation metabolite, 5-hydroxyindoleacetic acid. Interestingly, KYNA in the CB inversely correlated with 5-hydroxyindoleacetic acid in the PFC.

**Conclusions:** thus, this work found alterations in KP in two brain areas belonging to the cortico-cerebellar-thalamic-cortical circuit associated with SZ symptomatology, with a possible impact across areas in 5-HT degradation.

## BACKGROUND

Schizophrenia (SZ) is a multifaceted psychiatric disorder in which dysfunction of particular neuronal circuits occurs. The cortico-cerebellar-thalamic-cortical circuit may contribute to negative symptomatology and cognitive deficits in SZ [1, 2]. The cerebellum (CB) sends multiple inputs to the prefrontal cortex (PFC) through the thalamus, as well as dopaminergic inputs through the ventral tegmental area (VTA). There are feedback loops from cortex to CB regulating the activity of PFC neurons. Thus, the prefrontal cortex is the end route of this circuit responsible of cognitive and negative symptoms in schizophrenia, while the cerebellum is a key area in this circuit integrating and modulating multiple inputs from cortical areas and be the responsible of sending output signals back to the same cortical region through the thalamus to correct errors in cortical region. Recently, modulation of cerebellar activity has been proposed as a novel therapeutic intervention in SZ [3]. The control of PFC activity through cerebellar circuits improved symptoms, being of particular relevance for the negative ones [4]. However, the molecular basis underlying this improvement remains unknown.

Additionally, cognitive impairments have been particularly ascribed to the kynurenine pathway (KP) of tryptophan (Trp) metabolism [5]. Trp is commonly thought to be a precursor of the neurotransmitter 5-HT, but it can also be degraded to kynurenine (Kyn) [6] by one of these three enzymes, Tryptophan 2,3-dioxygenase (TDO), indoleamine 2,3-dioxygenase 1 (IDO1), and the less studied indoleamine 2,3-dioxygenase 2 (IDO2) [7]. Next, the Kyn metabolism trifurcates into distinct branches. It can be converted by kynurenine aminotransferases (KATs I-IV), mostly by KATII, to Kynurenine Acid (KYNA). It can also be oxidized by kynurenine 3-monooxygenase (KMO) to produce 3-hydroxykynurenine (3HK). Lastly, Kyn can undergo oxidative cleavage by Kynureniase (KYNU) to form anthranilic acid (AA). Both 3HK and AA leads to the formation of quinolinic acid (QA) [8].

KYNA and QA are neuroactive compounds with purported links to neuropsychiatric diseases [9]. QA is an N-methyl-D-aspartate receptor (NMDAR) agonist that can cause excitotoxicity, whereas KYNA is an NMDAR antagonist that can protect against excitotoxicity and apoptosis [10]. The KP in brain is compartmentalized between different cellular types. IDOs are expressed in neurons, astrocytes and microglia, TDO is highly expressed in astrocytes and in some neurons, and KMO is predominantly expressed in microglia [11]. Thus, some authors have simplified the scenario, suggesting that astrocytes produce KYNA in the “hypoglutamatergic branch” of the KP, and microglia generates QA, responsible for the “neurotoxic branch” [11]. In SZ, increased brain KYNA may lead to excessive NMDAR blockade, which could trigger psychotic symptoms and cognitive deficits [12]. Apart from glutamate, these neuroactive metabolites can modulate other systems of neurotransmission, particularly those dependent on monoamines, such as dopamine (DA) and 5-HT [13]. Both neurotransmitters have been implicated in SZ symptoms [14, 15]. Specifically, an excess of DA in subcortical areas and 5-HT hyperactivity at the serotonin 2A receptors (5-HT2A) on glutamate neurons in the cerebral cortex contribute to positive symptoms, while hypodopaminergic activity in the PFC is thought to cause negative symptoms [16, 17].

All of the elements of the KP are formed in the brain and in periphery [18] and regulated by pro-inflammatory cytokines, such as interleukin 1β (IL-1β), interferon-γ and tumour necrosis factor α, which are considered main actors of neuro-immune interactions [19]. Moreover, the study of anti-inflammatory cytokines such as IL-4 and IL-10 is growing, due to the current evidence supporting the existence of a dysregulated pro/anti-inflammatory cytokine balance in SZ [20]. In particular, IL-10 levels were decreased in postmortem brain PFC samples of SZ patients [21].

The dysregulation of the cytokines balance in SZ is associated to alterations on KP [22], and in particular, with the increased levels of KYNA observed at central level [23]. However, the precise relationship between inflammation and KP, and whether these interactions could be implicated in the symptomatology of SZ remain partly unknown.

The aims of this study were: 1) to investigate the possible alterations of the KP in brain PFC and CB in chronic SZ patients; 2) to explore the possible regulation of KP by pro/antiinflammatory cytokines; and 3) to search for correlations between KP and monoamine metabolic pathways, and SZ symptomatology, in the last term.

## METHODS

### Post-mortem human brain samples

Post-mortem human brain samples from the dorsolateral PFC (Brodmann area 9) and the cerebellum (CB; lateral cerebellar cortex) of patients with chronic schizophrenia (SZ; PCF, n =15; CB, n = 15) and of control subjects with no history of psychiatric disorders (PFC, n=14; CB, n=13) were obtained from the collection of Neurological Tissue of Sant Joan de Déu (SZ samples) [24] and the Brain Bank at the Hospital Universitari de Bellvitge (Control samples). CB and PFC samples were matched from the same subjects except from one control in each cohort. The demographic and clinical characteristics for these SZ subjects have been previously reported [25] and are described in detail in Table S1 in supplemental information. Possible tardive dyskinesia side effect of treatments was assessed in donors using the Abnormal Involuntary Movement Scale (AIMS)[26]. Total score was calculated using the sum of items 1 to 7, which assess the severity of abnormal movements in different regions of the body. Each item was classified from 0 to 4 according to the severity (0-absence, 1-minimum, 2-mild, 3-moderate, 4-severe). The mean and standard deviation of AIMS total score was of 5.05 ± 7.06 (n=16) with a minimum value of 0 and a maximum value of 23.50% (n=8) of the subjects showed absent symptoms. 6.25% (n=1) of the subjects showed minimum severity of symptoms, 31.25% (n=5) mild, 6.25 % (n=1) moderate and 6.25% severe (n=1). No subjects showed severe symptoms.

The study was approved by the Institutional Ethics Committee of Parc Sanitari Sant Joan de Déu. Written informed consent was obtained from each subject.

Specimens, extending from the pial surface to white matter and only including grey matter, were dissected and stored at −80 °C.

### Preparation of biological samples

Tissue samples were homogenized by sonication in phosphate-buffered saline (PBS) mixed with a protease inhibitor cocktail (Complete^®^, Roche, Spain) (pH =7), followed by centrifugation at 12,000 g for 10 min at 4°C.

### Cytokines and chemokine receptor levels

IL-1β (Raybiotech, ELH-IL1b), IL-4 (Cusabio, CSB-E04633h) and IL-10 (Raybiotech, ELH-IL10) tissue levels were detected using a commercially ELISA-based kit following the manufacturer’s instructions. For IL1β, the detection range is: 0.3 pg/ml −100 pg/ml, for IL10: 1 pg/ml - 150 pg/ml, and for IL4: 6.25 pg/ml-400 pg/ml. The intra- and interassay CV were <10% and <12% for IL1β and IL10; and <8% for IL4. Control values in PFC samples were: 0.59 +−0.12 (SD) pg/mg of tissue for IL1β, 0.32+−0.06 pg/mg of tissue for IL-10 and 7.34+−1.19 pg/mg of tissue for IL4. Control values in CB samples were: 0.57 +−0.06 (SD) pg/mg of tissue for IL1β, 0.47+−0.06 pg/mg of tissue for IL-10 and 8.44+−1 pg/mg of tissue for IL4.

### Western blot analysis

After determining and adjusting protein levels, homogenates were mixed with Laemmli sample buffer (Bio-Rad, USA), and 15 μg were loaded into an electrophoresis gel. Once separated on the basis of molecular weight, proteins from the gels were blotted onto a nitrocellulose membrane using a semi-dry transfer system (Bio-Rad) and were incubated with specific primary antibodies against: IDO1, IDO2, TDO; KMO, KATII and β-actin.

Primary antibodies used: (1) Indoleamine 2,3-dioxygenase 1 (IDO1, 1:750 in BSA 1%; sc25809, SCT); (2) Indoleamine 2,3-dioxygenase 2 (IDO2, 1:750 in BSA 1%; NBP2-44174, Novusbio); (3) Tryptophan 2,3-dioxygenase 2 (TDO, 1:500 in BSA 1%; ab123403, Abcam); (4) Kynurenine 3-Monooxygenase (KMO, 1:750 in BSA 1%; NBP2-29936, Novusbio); (5) Kynurenine aminotransferase 2 (KATII, 1:1000; sc377158, SCT); (6) β-actin (1:10000; A5441, Sigma).

Primary antibodies were recognized by the respective horseradish peroxidase-linked secondary antibodies. Blots were imaged using an Odyssey Fc System (Li-COR, Biosciences, Germany) and were quantified by densitometry (NIH ImageJ software). In all the WB analyses, the housekeeping β actin was used as a loading control, and every western blot was performed at least three times in separate assays. The data were presented as percentage change from control group.

### Kynurenine pathway metabolites measurement

KYNA (Cloud-Clone, CED718Ge) and QA (Cloud-Clone, CEK552Ge) tissue levels were detected using a commercially ELISA-based kit following the manufacturer’s instructions. For KYNA, the detection range is: 2.47-200ng/ml, and for QA: 1.23-100ng/ml. The intra- and interassay CV were 10% and 12% for KYNA and QA. Control values in PFC samples were: 808.64+−76.3(SD) pg/mg of tissue for KYNA, and 2.74+−0.37 pg/mg of tissue for QA. Control values in CB samples were: 1155.81+−110.43(SD) pg/mg of tissue for KYNA, and 4.33+−0.7pg/mg of tissue for QA.

### Monoamines

All samples were prepared in the same conditions. One mg of tissue was homogenized in 9 μl of acidic buffer (2 mL of 0.8 mol/L perchloric acid, 40 μL EDTA 0.1 mmol/L and 6 mL of ultrapure H20) at 4°C and stored at −80°C until analysis. Then, after centrifugation (1500 g, 10 minutes, 4°C), the supernatant was recovered and mixed 1:2 (v/V) with the chromatographic mobile phase (0.1 mol/L citrate/acetate buffer, 1.2 mmol/L EDTA/heptanesulphonic acid dissolved in 7% methanol: final pH=4). The monoamines were measured by ion-pair HPLC coupled to electrochemical detection. This procedure was fully validated in our laboratory, as previously reported [27]. This procedure is accredited by the ISO 15189 norm (ENAC agency) and it is subjected to an external quality control scheme (www.ERNDIM.org).

### Network interaction analysis

Interaction between the elements of KP and its relationship with pro/anti-inflammatory cytokines and monoamine metabolites were evaluated using a co-expression analysis adapted from a previous study in SZ [28]. This co-expression analysis for characterising a pathway in a biological substrate is based on the assumption that proteins functionally related co-expressed [29]. Briefly, among all samples studied (control and SZ samples), only those with the highest (75th percentile) and lowest (25th percentile) fold-change values of the upstream member of the pathway were selected for analysis, starting from TDO, IDO1 and 2 (i.e. High-IDO1 group or Low-IDO1 group). Correlation analyses correlations with the downstream member of the pathway were performed. Differences in the protein expression levels of the downstream member between the group with samples with the highest protein levels and that with the lowest protein levels were evaluated. Significant correlations in this co-expression analysis indicate that the interaction distance between these two proteins is short in this network [30]. The pathway was “coupled” when all the downstream members of the pathway correlate with the upstream member, reflecting the proximity of the members in the pathway. The term “uncoupled pathway” was used when no significant correlations were detected between the downstream proteins and the upstream member in the pathway, indicating a higher distance between the elements of the pathway.

### Statistical analysis

Quantitative values for protein or activity were tested for Gaussian distribution using the D'Agostino & Pearson omnibus normality test and compared using Student's unpaired t-test or Mann-Whitney test according to the distribution of each variable. Outliers were detected using the ROUT method for parametric variables with a significance Q set at 1% for the detection and a Pierce test for the non-parametric variables. Spearman or Pearson correlation analyses were carried out according to the distribution of each variable to detect association of the molecular measures with symptom severity. The False Discovery Rate (FDR) assessed with the Benjamini and Hochberg method was computed for all p values resulting from comparisons with symptoms in each brain region and was set to 0.1. Spearman or Pearson correlation analyses were also performed to assess association of the molecular measures with and potential confounding factors (age, post-mortem delay, pH, chlorpromazine dose (CPZd) and duration of illness). In case of finding a significant association with a potential confounder factor, a multiple linear regression model with a stepwise forward procedure was used to adjust to the co-variable where indicated. Statistical analyses were performed with GraphPad^©^ Prism version 6.01 and SPSS 24. All statistical tests were two-tailed and the significance level was set to 0.05.

## RESULTS

### Expression of pro/anti-inflammatory cytokines and KP elements in PFC and CB. Correlations with SZ symptomatology

IL-10 protein expression was significantly decreased in SZ subjects compared to control in the PFC (Fig. 1A). We did not find differences in IL-1β or IL-4 between groups (Figs. 1A-B).

**Figure 1.**
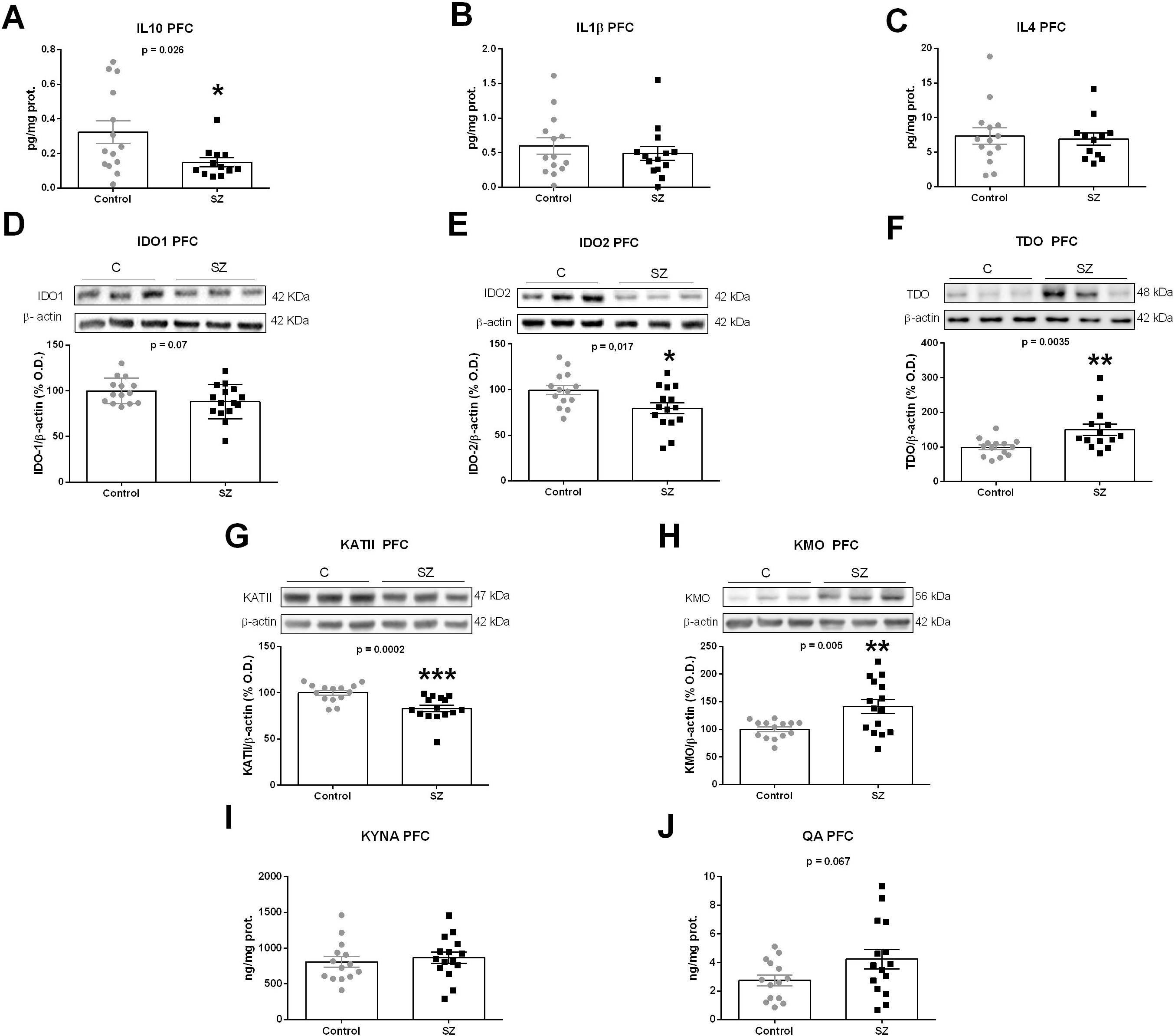
Kynurenine pathway in post-mortem brain PFC samples of subjects with chronic SZ and matched controls. IL-10 (A), IL-4 (B), IL-1β (ELISA) (C), IDO1 (D), IDO2 (E), TDO (F), KMO (G), KATII (H) (Western Blot), KYNA (I) and QA (J) (ELISA) levels in PFC of SZ subjects (black squares) and matched controls (grey circles). (B) The densitometric data of the respective bands of interest are normalized with respect to β-actin (lower band). The blots are representative of all data analyzed and have been cropped to improve the clarity and conciseness of the presentation. *p < 0.05; **p < 0.01; ***p < 0.001 vs. control (C). ^a^Student's unpaired t-test or ^b^Mann-Whitney test. Data represent the mean ± standard error of the mean (N=14 control and 15 SZ). IL-10: interleukin-10; IL-4: interleukin-4 IL-1β: interleukin 1 Beta; IDO1: indoleamine 2,3-dioxygenase 1; IDO2: indoleamine 2,3-dioxygenase 2; TDO: Tryptophan 2,3-dioxygenase 2; KMO: kynurenine monooxygenase; KATII: kynurenine aminotransferases II.

We explored the branch of the KP composed by IDO-1 and 2 and KMO generating QA, and the alternative branch composed by TDO and KATII that lead to KYNA production. The respective expression of both IDO enzymes decreased in the PFC of SZ subjects compared to control, especially for IDO-2 (Figs. 1D-E). TDO was increased in SZ while KAT II was markedly decreased (Figs. 1F-G) Contrarily, the expression of KMO was increased in SZ (Fig. 1H). Accordantly, QA tended to increase in SZ (p=0.07) (Fig. 1J). However, no differences were found for KYNA content in the PFC (Fig. 1I).

Furthermore, we analysed the possible influence of other variables on the enzymes that significantly changed between groups (Table S2). The only significant correlation was found between TDO and post-mortem delay (PMD) (Table S2). Linear regression analysis revealed that the increase of TDO in SZ remained significant after adjusting by PMD (β =0.456, p =0.015, adjusted R2 0.244).

We then explored whether KYNA and QA levels in the PFC correlate with the severity of the symptoms. KYNA did not correlated with negative symptoms in SZ patients (r Pearson =0.520, uncorrected p=0.046, FDR-adjusted p=0.188).

The same inflammatory and KP-related parameters were measured in the CB. There were no significant changes in any of the mediators measured (Fig. 2A-J).

**Figure 2.**
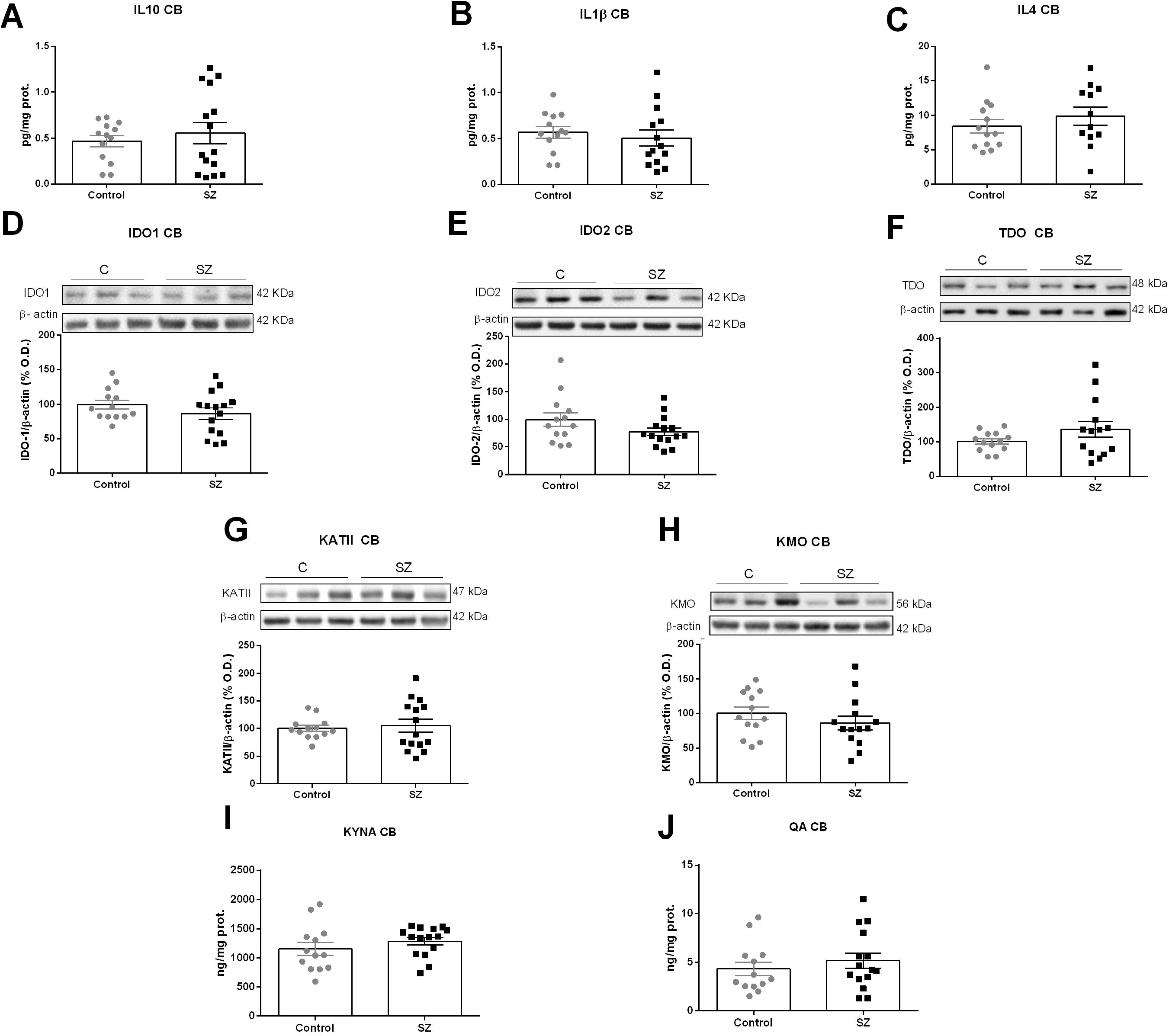
Kynurenine pathway in post-mortem brain CB samples of subjects with chronic SZ and matched controls. IL-10 (A), IL-4 (B), IL-1β (ELISA) (C), IDO1 (D), IDO2 (E), TDO (F), KMO (G), KATII (H) (Western Blot), KYNA (I) and QA (J) (ELISA) levels in CB of SZ subjects (black squares) and matched controls (grey circles). (B) The densitometric data of the respective bands of interest are normalized with respect to β-actin (lower band). The blots are representative of all data analysed and have been cropped to improve the clarity and conciseness of the presentation. Data represent the mean ± standard error of the mean. (N=13 control and 15 SZ). IL-10: interleukin-10; IL-4: interleukin-4 IL-1β: interleukin 1 Beta; IDO1: indoleamine 2,3-dioxygenase 1; IDO2: indoleamine 2,3-dioxygenase 2; TDO: Tryptophan 2,3-dioxygenase; KMO: kynurenine monooxygenase; KATII: kynurenine aminotransferases II.

We also analysed whether KYNA and QA levels in the CB, respectively, correlate with the severity of the symptoms. Contrarily to the PFC, KYNA in the CB inversely correlated with both negative (r Pearson=−0.734, uncorrected p=0.002, FDR-adjusted p=0.008) and general psychopathological subscales (r Pearson=−0.692, uncorrected p=0.004, FDR-adjusted p=0.008) in SZ.

Moreover, we have analysed the possible influence of other variables on KYNA levels. Significant correlations were found between KYNA in the CB and CZPd.

A linear regression analysis revealed that the correlation between KYNA and both negative and general psychopathological PANSS remained significant after adjusting by CPZd (KYNA-Negative PANSS: β= −1.157, p = 0.001, adjusted R2=0.662; KYNA-General PANSS: β = −0.787, p=0.032, adjusted R2=0.485).

### Network interaction analysis of the KP and cytokines

A co-expression analysis (Figures 3A-B) showed an inverse correlation between IL-10 and TDO in the PFC (r Pearson=−0.801, p value=0.003). Moreover, interactions between proteins showed significant associations between TDO and KMO (r Pearson=0.692, p value=0.009), and KMO and QA (r Spearman=0.565, p value=0.035) (Fig. 3B). A comparison between the groups showed increased levels of KMO in individuals with higher levels of TDO compared to individuals with reduced TDO (TDO-KMO: t test, p value=0.021) (Fig. 3C). A tendency to associate is present between KMO and QA (KMO-QA: t test, p value=0.06) (Fig. 3C). These results indicate that IL-10 is acting upstream of TDO with the coupled TDO-KMO-QA pathway, which was altered in the PFC.

**Figure 3:**
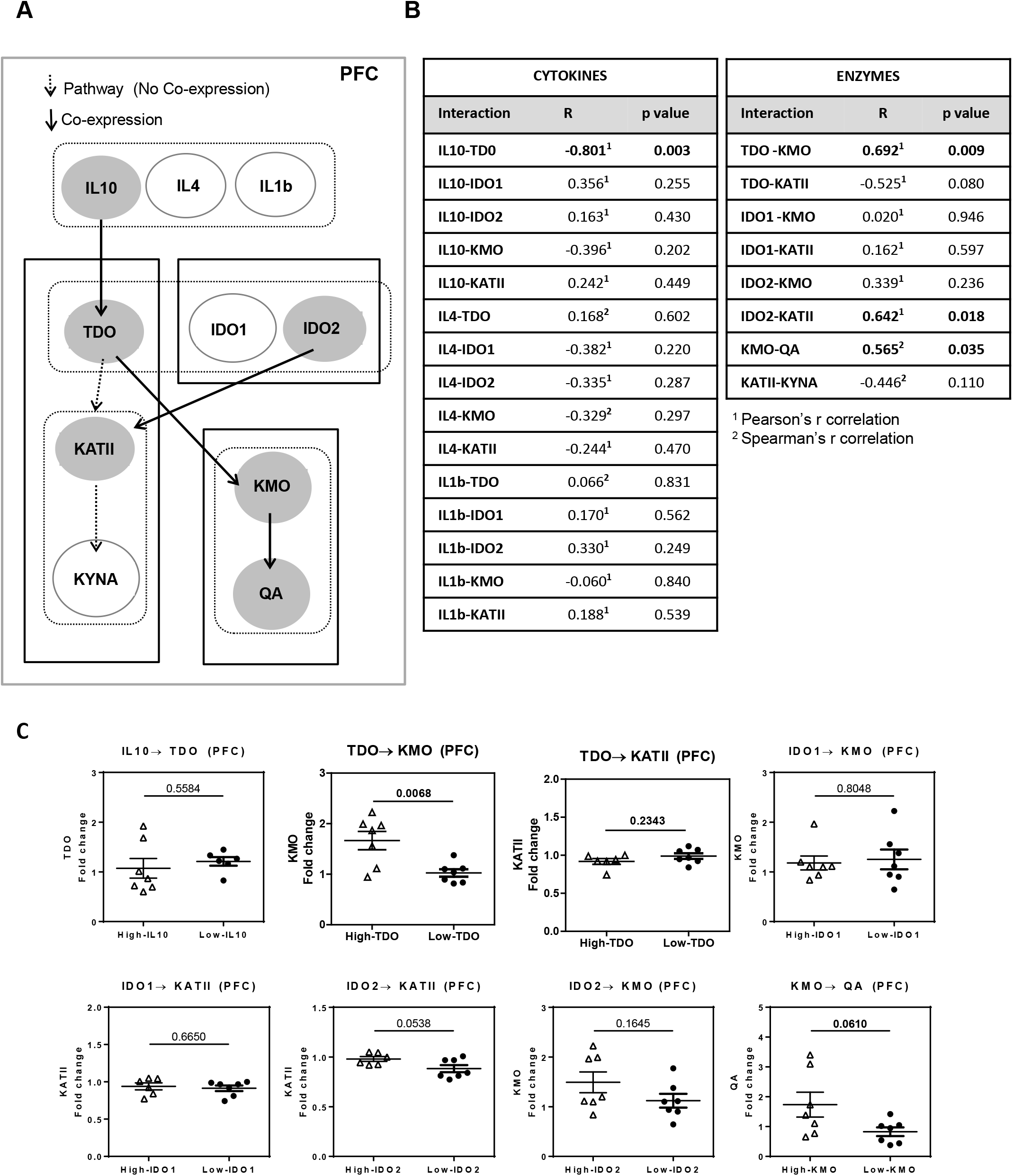
Co-expression analysis of Kynurenine pathway in postmortem prefrontal cortex. (A) Network model of Kynurenine pathway in the postmortem prefrontal cortex. Samples with the highest and lowest expression levels were selected for the analysis. The white arrows show signal transduction between upstream and downstream elements. Black arrows indicate significant co-expression with the downstream element. Filled circles indicate a significant correlated or changed element in interaction analysis (p<0.05). (B) Correlations using Pearson or Spearman analysis of the samples with the highest expression and lowest expression of upstream elements among all 29 samples were selected. Significant interactions are indicated with bold (p<0.05). (C) Levels of the downstream element were compared between High- and Low-groups of the upstream element by unpaired t test or Mann-Whitney test (p<0.05). Data represent the mean ± standard error of the mean.

Our analysis also showed an interaction between IDO2 and KATII (r Pearson=0.642, p value=0.018).

In the CB, we found an inverse association of IL-1β with TDO (r Spearman=−0.836, p value= 0.001), and between TDO and KATII (r Pearson=−0.603, p value=0.038) (Figs. 4A-B). KATII is uncoupled with its metabolite KYNA. Individuals with lower levels of IL-1β showed increased expression of TDO (IL-1β-TDO: Mann Whitney U, p value=0.051), and individuals with lower levels of TDO also expressed increased levels of KATII (TDO-KATII: Mann Whitney U, p value=0.051) (Fig. 4C). Furthermore, we found that IL-10 interacted downstream in the pathway, at the level of KMO (r Spearman=−0.597, p value=0.031) (Fig. 4C). These results indicate that, contrary to the PFC, the TDO-KMO-QA pathway is uncoupled in the CB, with IL-10 acting on KMO.

**Figure 4:**
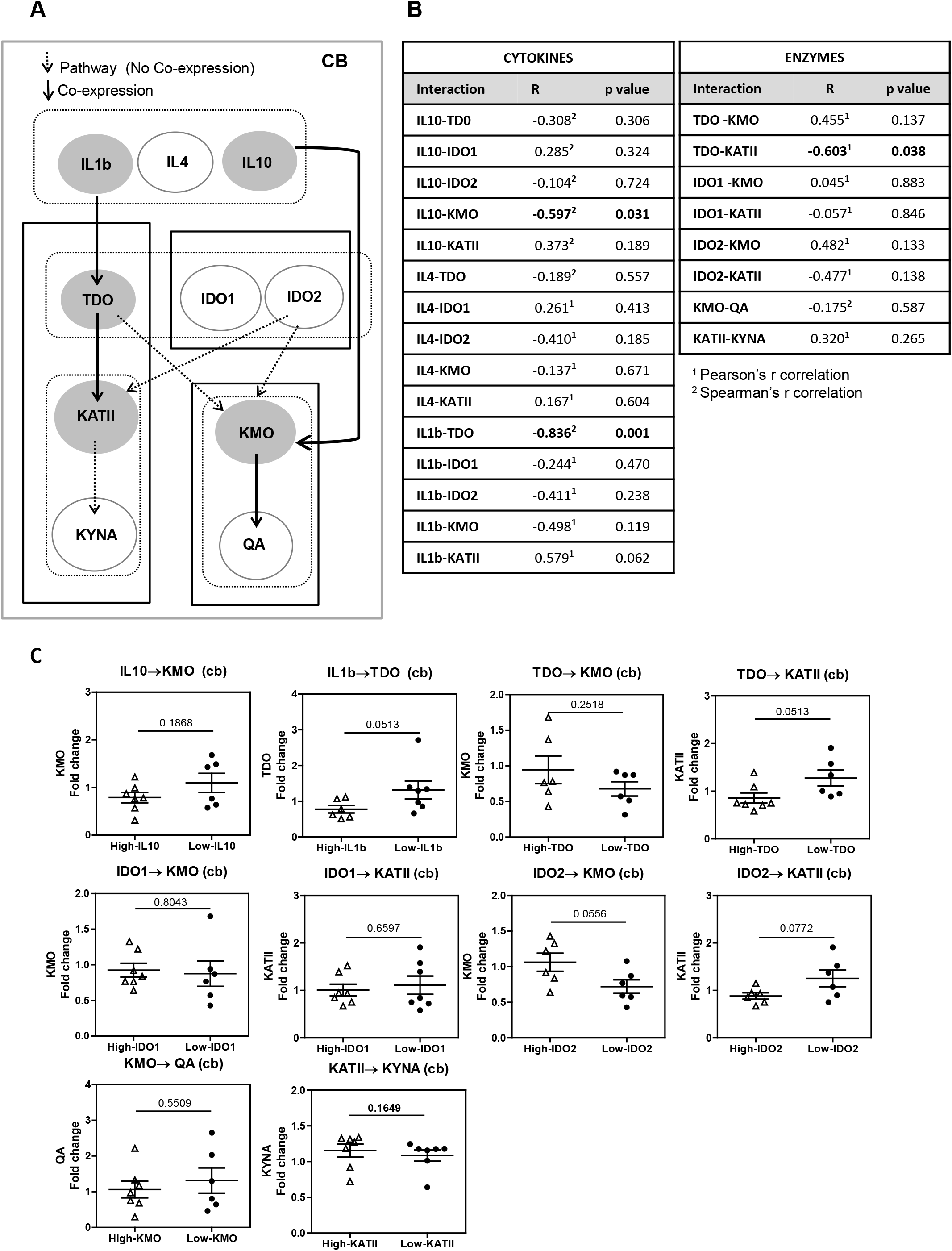
Co-expression analysis of Kynurenine pathway in postmortem cerebellum. (A) Network model of Kynurenine pathway in the postmortem cerebellum. Samples with the highest and lowest expression levels were selected for the analysis. The white arrows show signal transduction between upstream and downstream elements. Black arrows indicate significant co-expression with the downstream element. Filled circles indicate a significant correlated or changed element in interaction analysis (p<0.05). (B) Correlations using Pearson^1^ or Spearman^2^ analysis of the samples with the highest expression and lowest expression of upstream elements among all 28 samples were selected. Significant interactions are indicated with bold (p<0.05). (C) Levels of the downstream element were compared between High- and Low-groups of the upstream element by unpaired t test or Mann-Whitney test (p<0.05). Data represent the mean ± standard error of the mean.

### Synthesis and metabolism of monoamines in PFC. Correlations with SZ symptomatology

DA and 5-HT systems in the PFC were studied by measuring the levels of DA and homovanillic acid (HVA), and 5-hydroxyindoleacetic acid (5HIAA), respectively. Other metabolites such as 3-orthomethyldopa (3-OMD) and methoxyhydroxyphenylglycol (MHPG) for the DA pathway, and 5-hydroxytryptophan (5HTP) for the 5-HT pathway were also tested [27].

There were no significant changes in any of the metabolites between groups (Fig. 5A). Another network analysis revealed that DA levels inversely associated with MHPG from noradrenergic neurons (r Pearson=−0.657, p=0.011), and showed a tendency to associate with HVA (Fig. 5B) (r Pearson=−0.525, p=0.054), suggesting that DA degradation pathways are coupled in the PFC. No other associations were found (Fig. 5B).

**Figure 5:**
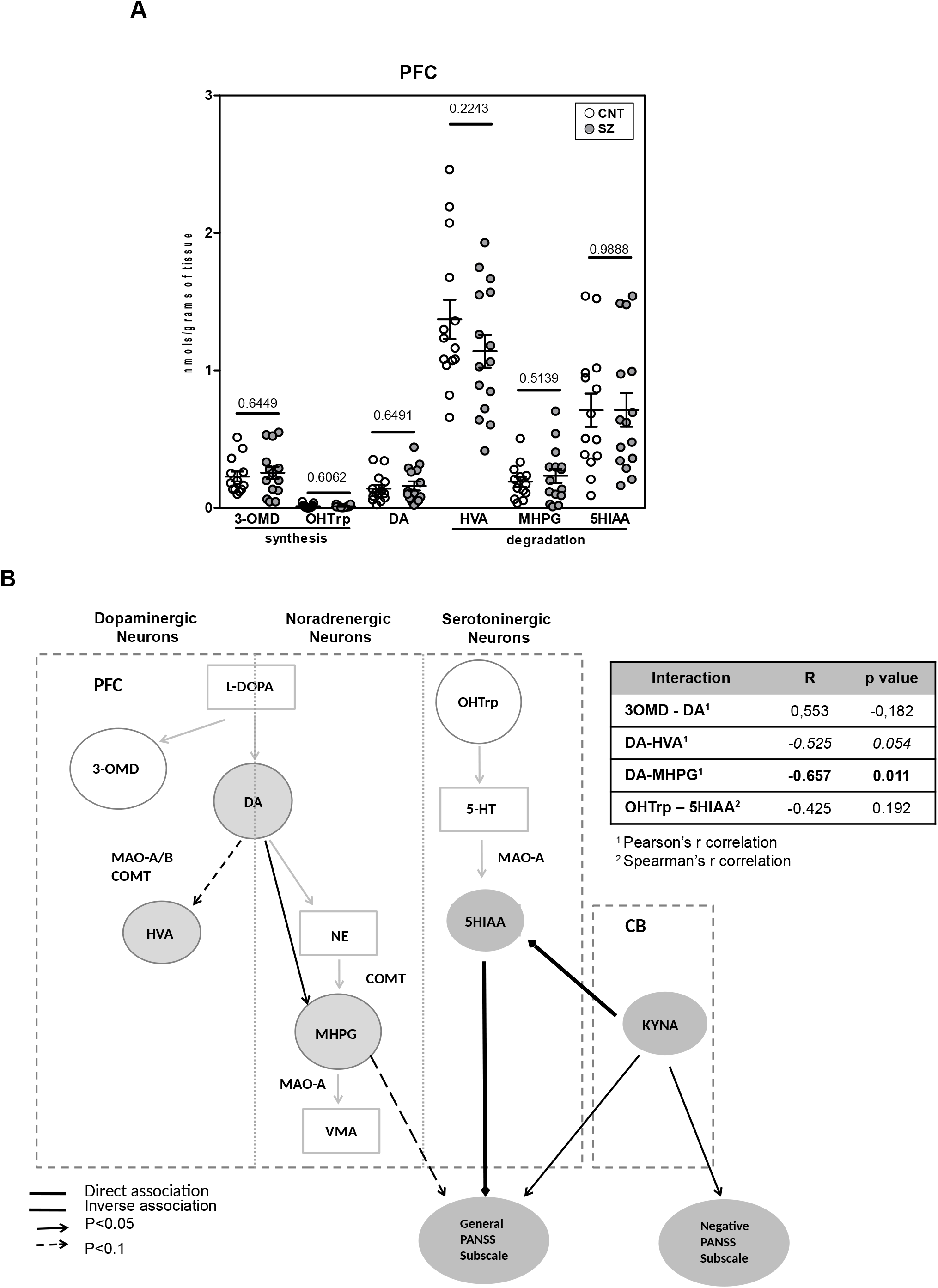
Correlations of the kynurenine pathway with Monoamine pathway and SZ symptomatology. (A) Differences in monoamine pathway elements between control (n=14) and schizophrenia group (n=15) in the postmortem prefrontal cortex by Ultra-High-Performance Liquid Chromatography-Tandem Mass Spectrometry. One outlier was detected for 3-OMD, MHPG and 3OHTrp in the control group and one in the schizophrenia group for 3-OHTrp and therefore excluded from the analysis. Results were expressed as pmol/gram of tissue. Statistical analysis was performed using unpaired t test or Mann-Whitney test (p<0.05). (B) Schematic representation of the significant interactions between dopaminergic, noradrenergic and serotoninergic pathway elements in the schizophrenia group (n=15) and final product elements from kynurenine pathway in the cerebellum (n=13) and prefrontal cortex (n=15). Analysis was carried out using Pearson^1^ or Spearman^2^ correlations. Correlations between elements of monoamine pathway are shown in the table. Significant associations are indicated in bold. The double line black arrows show an inverse association and the single line black arrows a direct association. between elements of the different pathways and general PANSS subscale scores (continued arrows p<0.05; discontinued arrows p<0.1).

Moreover, we explored the possible association between DA levels and SZ symptoms. No significant correlation was found. We also explored the possible correlations between the expression of the degradation metabolites of monoamines in PFC with SZ symptoms. We found a significant correlation between PANSS general psychopathological subscale and the metabolite 5HIAA (degraded by monoamine oxidase A (MAO-A)) (Pearson r=0.614, uncorrected p=0.015; FDR-adjusted p value=0.090; p value threshold=0.017) expression in PFC. A tendency to associate with PANSS general was also found for MHPG (substrate of MAO-A (Pearson r=−0.448, p=0.094). These results suggest a possible relationship between SZ symptoms and an increase activity of MAO-A in noradrenergic and serotoninergic neurons in the PFC.

### Correlations of the KP with monoamine pathway and SZ symptomatology

We further explored the existence of correlations between KYNA in the CB and 5HIAA and MHPG in the PFC. We found an inverse correlation between KYNA in CB and 5HIAA in PFC (r Spearman=−0.602, p=0.023). These results suggest a possible modulation of 5-HT degradation in the PFC mediated by KYNA inhibitory action on glutamatergic neurons in the CB (Fig. 5B).

## DISCUSSION

We have found alterations in the KP in the cortico-cerebellar-thalamic-cortical circuit with a possible impact across areas in the metabolism of 5-HT and associated to general psychopathological symptoms severity in SZ.

In our study the changes observed in KP are exclusive of the PFC. Previous studies also focused on PFC reporting an increase in TDO expression and activity in SZ patients [23]. IDOs has been less explored: although one previous study did not found changes at mRNA level [31], we have found a decreased protein expression in SZ. In the case of KATII and KMO our results are opposite to previous ones in which KATII expression was increased or unchanged [23, 32], and KMO was decreased [32]. These changes in KATII and KMO were found at mRNA level and in their enzymatic activity while we observed changes at the level of protein expression. Regarding KYNA and QA, we have only found a trend to increase in QA (p=0.07), while others have focused on KYNA, reporting a consistent increase in brain in SZ, PFC included [23, 33].

Our findings of increased expression of TDO in SZ in PFC suggest that the conversion of Trp to kynurenine is promoted probably in astrocytes. Moreover, the fact that TDO is coupled with increased KMO in SZ in PFC, suggests a putative crosstalk between both cellular types. This raises the possibility that in SZ, kynurenine is produced by astrocytes, released to the extracellular fluid and transported into microglia cells through LAT1 or the Leucine/Arginine transporter Slc7a7 [34, 35]. This mechanism could compensate the decreased levels of IDO2 detected in PFC in our study, enhancing the production of “neurotoxic” QA by microglia in PFC. However, further studies are needed to explore the precise cellular types and associations involved.

There are no previous data for KP alterations in CB postmortem samples in SZ to compare with our negative results, but the existence of functional alterations or in the expression at mRNA level/activity of the different elements of KP need to be further explored.

The network interaction analysis made allowed us to explore possible interactions between the different elements of the KP and their regulatory mechanisms above their net expression levels. Thus, our analysis suggested the role of IL-10 as an endogenous regulator of the KP in PFC and CB, although at a different level. Decreased protein and mRNA expression of IL-10 in SZ has been also reported previously at mRNA level in PFC, as well as no differences in IL-1β, as in our experimental conditions [21, 30].

In our study, IL-10 is inversely related to the expression of TDO in PFC and KMO in CB, respectively. A former study showed pro-inflammatory effects of QA via significant decreases of IL-10 at systemic level in healthy volunteers [36]. The opposite relationship is also possible. Thus, IL-10 derived from T lymphocytes decreased IDO1 expression in the resolution phase of inflammation in the PFC of mice [37]. In our conditions, it remains to be elucidated whether the decreased levels of IL-10 in PFC observed are a consequence of the up-regulation of coupled TDO to KMO and subsequent QA generation. In CB, IL-10 is also inversely associated with this microglial pathway, being coupled directly to KMO, and not related to TDO. No effect on QA was observed in CB. The selective inhibition of microglial KMO is an emerging neuroprotective target [38].

Although we have not find changes in the expression levels of IL-1β in the two brain areas studied, the network interaction analysis in CB showed that IL-1β was inversely related to TDO (p=0.001) and directly related with KATII, although in a lesser degree (p=0.06). In our knowledge there are no data about the relationship between IL-1β and KATII, but IL-1β has potential to induce KYNA in human astrocyte cultures increasing TDO expression, and also a strong correlation between IL-1β and TDO2 expression in human brain post-mortem prefrontal grey matter from healthy donors was also shown [39]. Whether the discrepancies observed are due to the chronic nature of SZ in a possible compensatory mechanism between IL-1β and TDO remains to be elucidated, but it is worth to remark that the pathway IL-1β-TDO-KATII is uncoupled to its final metabolite KYNA in CB, being the levels of KYNA similar to control samples.

Our correlation analysis point to KYNA as the main element of KP that correlated with SZ symptomatology. As commented above, previous evidences showed increased KYNA levels in SZ and in animal models of the disease [23, 32, 33, 40]. This increase has been related to SZ symptomatology (mainly in cognition) by means of alterations in glutamatergic, acetylcholinergic, GABAergic and dopaminergic neurotransmission [41–44]. However, there are no data about KYNA levels in CB, and their relation with SZ symptoms remain unexplored. It is worth to mention that some authors have found reduced levels of KYNA in the plasma of SZ patients [45]. Thus, all the previous evidence suggests that KYNA levels in SZ could vary between brain and periphery, and in the different states and clinical subtypes of the disease, being affected by diverse confounding factors, such as the severity of inflammation or antipsychotics [33].

Although changes in monoamine degradation metabolites have been reported in CSF and plasma in SZ [46], our analysis did not reveal any change. Only a few studies have focused on monoamine metabolism in postmortem brain, reporting an increase on DA or HVA in subcortical areas in SZ [47]. However, to our knowledge, no data were available for PFC in SZ.

There is an emerging interest of how modulating cerebellar activity induces changes in symptoms, being proposed that cortico-cerebellar-thalamic-cortical and the cerebellar-VTA-cortical circuits may underlie this effect by regulating PFC activity [3, 4]. We further explored the possibility that the levels of KYNA in CB could be related to monoamine metabolism in the PFC in the context of symptom severity in SZ. Since a glutamatergic/dopaminergic circuit has been recently identified between CB and PFC through the VTA, the most plausible hypothesis to explain cerebellar KYNA-related negative symptoms (associated with a hypodopaminergic activity in PFC) was effects on DA metabolites. Surprisingly, either DA nor HVA in PFC correlated to KYNA cerebellar levels and negative symptoms.

However, we found a correlation between levels of KYNA in CB and 5-HIAA in PFC linked to general psychopathology symptoms. A trend to be associated with this subscale was also detected for MHPG. 5-HIAA and MHPG are the product and the substrate of the same enzyme, the Monoamine Oxidase A (MAO-A), respectively. This data raises the possibility that there may be increased MAO-A activity in serotoninergic and noradrenergic neurons in the PFC in subjects with more severe general psychopathological symptoms (affective, excitement and negative). We also found a direct interaction between HVA and 5-HIAA that agrees with the idea of increased MAO-A activity in PFC in SZ. Unfortunately, there are no previous studies exploring the activity of MAO-A in postmortem brain PFC in SZ. This possible increased activity could result in a faster removal of 5-HT and noradrenaline neurotransmitters in this area linked to severity in symptoms. This is in agreement with the proposed reduction of 5-HT in brain in SZ responsible for emotional and affective symptoms [14]. However, there is not a consensus and other authors proposed an increase of 5-HT release, a hyperactive effect on 5HT2A receptors on glutamatergic neurons in cerebral cortex and a global higher serotoninergic tone in SZ [15].

5-HIAA and MHPG levels in plasma in SZ have been inversely and directly associated with depression/anxiety or excitement components of PANSS scale, respectively [48], although some authors have alerted about the difficulty to correlate the levels of blood/urine monoamines with brain neurotransmission status, even in severe genetic conditions affecting DA and 5-HT metabolism [49]. However, the associations detected for these two metabolites with PANSS symptoms in our study are in the opposite direction. In agreement with our findings, 5-HIAA in CSF has also been reported to positively associate with several symptoms including some of the general psychopathological subscale [46]. In addition, previous studies has reported positive associations between 5-HIAA and KYNA in CSF and plasma in first episode psychosis, which are the result of a global metabolism of different brain areas [50]. However, our findings showed an inverse correlation between KYNA levels in CB and 5-HIAA levels in the PFC suggesting a more complex interrelation between both brain areas. This connection might be mediated by glutamatergic outputs from CB that could be impacting on 5-HT degradation in the PFC. However, the neural bases of the implicated circuits are not clear.

Our study has some limitations: firstly, our cohort consisted of males only and it is needed to investigate possible gender differences. Second, although symptoms and in particular, negative symptoms, may be stable measures in chronic SZ patients [51], small variations in the clinical scores up to death may still slightly affect the associations we report here. Third, the patients with chronic SZ had long-term and heterogeneous antipsychotic medications. We have controlled the possible effect of antipsychotics on our findings using CPZd. This allowed us to identify an effect of antipsychotic treatment on KYNA levels in the CB. Fifth, the co-expression analysis is a prediction of how members of KP are functionally related among them and with monoamine metabolites. Future experimental interaction analyses with larger cohorts are needed to confirm these results.

## DECLARATIONS

## Consent for publication

Not applicable

## Acknowledgements

Not applicable

## Availability of data and materials

The datasets used and/or analysed during the current study are available from the corresponding author on reasonable request.

## Authors’ contributions

Amira Ben Afia, Elia Vila and Aida Ormazabal contributed to the acquisition and analysis of the results; Karina MacDowell, Juan Carlos Leza, Josep Maria Haro and Rafa Artuch contributed to the interpretation of data for the work and revised critically for important intellectual content; and Belen Ramos and Borja García-Bueno designed and drafted the work.

