## Supplemental Material for "Kynurenine pathway in post-mortem prefrontal cortex and cerebellum in schizophrenia: relationship with monoamines and symptomatology"

**SUPPLEMENTAL INFORMATION**

**Material and Methods**

Post-mortem human brain samples

Specimens, extending from the pial surface to white matter and only including grey matter, were dissected and stored at −80 °C.

**Table S1: Demographic, clinical and tissue-related features of cases**

|  | Chronic Schizophrenia | Non-psychiatric control | Statistic | p value |
| --- | --- | --- | --- | --- |
| *Prefrontal cortex* | (n=15) | (n=14) |  |  |
| Gender | Male- 100% (n=15) | Male- 100% (n=14) | N/A | N/A |
| Age at death | 74 ± 9 years | 74 ± 9 years | 97.5^a^ | 0.760 |
| PMD | 4.9 ± 2.2 hours | 5.7 ± 1.6 hours | 1.08; 27^b^ | 0.290 |
| pH | 6.6 ± 0.4 | 6.9 ± 0.4 | 74.00^a^ | 0.173 |
| Age of onset of illness | 22 ± 7 years | N/A | N/A | N/A |
| Duration of illness | 52 ± 10 years | N/A | N/A | N/A |
| D-A interval^c^ | 19 ± 14 months | N/A | N/A | N/A |
| Daily AP dose^d^ | 589 ± 491 mg/day | N/A | N/A | N/A |
| Clinical Scales |  | N/A | N/A | N/A |
| PANSS Positive | 24.4 ± 7.7 | N/A | N/A | N/A |
| PANSS Negative | 31.9 ± 10.6 | N/A | N/A | N/A |
| PANSS General | 51.7 ± 15.3 | N/A | N/A | N/A |
| *Cerebellum* | (n=15) | (n=13) |  |  |
| Gender | Male- 100% (n=15) | Male- 100% (n=13) | N/A | N/A |
| Age at death | 75 ± 10 years | 74 ± 9 years | 0.23; 26 ^b^ | 0.816 |
| PMD | 4.6 ± 2.5 hours | 5.9 ± 1.6 hours | 1,63; 26^b^ | 0.115 |
| pH | 6.5 ± 0.4 | 6.8 ± 0.6 | 70.5 ^a^ | 0.212 |
| Age of onset of illness | 22 ± 7 years | N/A | N/A | N/A |
| Duration of illness | 52 ± 10 years | N/A | N/A | N/A |
| D-A interval^c^ | 19 ± 13 months | N/A | N/A | N/A |
| Daily AP dose^d^ | 589± 491 mg/day | N/A | N/A | N/A |
| Clinical Scales |  | N/A | N/A | N/A |
| PANSS Positive | 25.1 ± 8.3 | N/A | N/A | N/A |
| PANSS Negative | 30.9 ± 9.6 | N/A | N/A | N/A |
| PANSS General | 53.3 ± 15.6 | N/A | N/A | N/A |

Mean ± standard deviation or relative frequency are shown for each variable; PMD, postmortem delay; D-A, Death to clinical assessment interval; AP, antipsychotic; PANSS, Positive and Negative Syndrome Scale; N/A, not applicable. The same individuals were included for both brain areas with the following exceptions in the SZ group: prefrontal cortex tissue was only available for one individual and cerebellum was only available for another individual due to the lack of sufficient tissue from these individuals.

^a^Mann-Whitney U is shown for non-parametric variables.

^b^T-statistic and degrees of freedom are shown for parametric variables.

^c^D-A Interval is defined as the number of months between clinical assessments and the time of death.

^d^Last chlorpromazine equivalent dose was calculated based on the electronic records of drug prescriptions of the patients.

Two patients were being medicated with first-generation antipsychotics, nine were medicated with second-generation antipsychotics, three were treated with a combination of first and second generation antipsychotics and two patients were antipsychotic-free. Of these, nine patients were treated with benzodiazepines, three were under biperiden treatment and one under mirtazapine treatment. In particular, the following antipsychotics were present: haloperidol (n=4), levomepromazine (n=2), quetiapine (n=4), olanzapine (n=7), risperidone (n=1), clotiapine (n=1), amisulpride (n=1).

**Results**

**Table 2: Association analysis of other variables in the SZ-C cohort.**

| **SZ-C cohort** | Age | PMD | pH |
| --- | --- | --- | --- |
| *Prefrontal cortex (n=29)* |  |  |  |
| IL10 | -0,275^2^ | 0,228^2^ | 0,151^2^ |
| IDO2 | -0.126^1^ | 0,283^1^ | -0,130^2^ |
| TDO | 0,135^2^ | **0,398^2,a^** | 0,170^2^ |
| KATII | -0.102^1^ | 0,231^1^ | 0,246^2^ |
| KMO | -0.187^1^ | -0,006^1^ | -0,295^2^ |

SZ, schizophrenia; C, control; PMD, *postmortem* delay; Significant associations are indicated in

bold ^a^p<0.05.

^1^ Pearson’s r is shown for parametric variables.

^2^ Spearman’s r is shown for non-parametric variables.
